# Proof of concept continuous event logging in living cells

**DOI:** 10.1101/225151

**Authors:** Andrey Shur, Richard M. Murray

## Abstract

Cells must detect and respond to molecular events such as the presence or absence of specific small molecules. To accomplish this, cells have evolved methods to measure the presence and concentration of these small molecules in their environment and enact changes in gene expression or behavior. However, cells don’t usually change their DNA in response to such outside stimuli. In this work, we have engineered a genetic circuit that can enact specific and controlled genetic changes in response to changing small molecule concentrations. Known DNA sequences can be repeatedly integrated into a genomic array such that their identity and order encodes information about past small molecule concentrations that the cell has experienced. To accomplish this, we use catalytically inactive CRISPR-Cas9 (dCas9) to bind to and block attachment sites for the integrase Bxb1. Therefore, through the co-expression of dCas9 and guide RNA, Bxb1 can be directed to integrate one of two engineered plasmids, which correspond to two orthogonal small molecule inducers that can be recorded with this system. We identified the optimal location of guide RNA binding to the Bxb1 attP integrase attachment site, and characterized the detection limits of the system by measuring the minimal small molecule concentration and shortest induction time necessary to produce measurable differences in array composition as read out by Oxford Nanopore long read sequencing technology.

## Introduction

Living cells are capable of detecting and responding to sophisticated stimuli present in their environment. Light [1], heat [2], chemicals, and proteins [3] represent a few of the types of stimuli that cells can distinguish. The ability to detect these stimuli is useful to the cell’s survival if the cell can appropriately respond to the stimulus; producing a heat-shock protein in response to heat, or a toxin efflux pump in response to toxins, for example. Many of these responses are short lived, such as a temporary increases in protein transcription or translation rate. For a bacterium, temporary changes are good because they represent a return to homeostasis. In many situations, however, it is useful to know the order and identity of these changes. A biologist must build sophisticated instruments to measure and record the identity of these stimuli using separate, macroscopic detection and recording devices. In this work, we describe a genetically encoded system that can create a record of chemical stimuli a cell has seen within that cell’s DNA.

Event recording can be divided into two modes of operation. First, the event is detected, and then it is recorded. As mentioned above, living cells naturally respond to stimuli in ephemeral ways, and much work has been done to characterize the signal transduction networks behind these responses. Transcriptional upregulation is an extremely common mechanism of response to stimuli in all living cells. Temporary, rapid transcriptional activation is enacted in bacteria through repressors unbinding promoters, or activators binding upstream of promoters and recruiting polymerase. In this work we make use of known promoters with well characterized activating/repressing proteins. The challenge, then, is how to convert this temporary transcriptional activation to a permanent genetic record. Previous work on DNA-based event recorders has focused on using phage integrases to irreversibly “flip” pieces of DNA in response to stimuli [4]–[6]. A temporary increase in RNA transcription, then, is converted through the action of integrase proteins into a permanent, lasting change in DNA. Phage integrases are extremely useful proteins to employ in this regard because their action to recombine specific DNA sequences is deterministic, fast [7], and irreversible. However, integrase-based event recorders that flip DNA typically have a limited number of attainable DNA states, meaning that only a few events can be recorded before the memory capacity is ‘used up’. Previous work using cas9 to stochastically excise memory units (conceptually similar to flipping memory units with integrases) in mouse stem cells[8] has shown that a modest number of memory units can be used to trace back cellular lineage over many generations. However, lineage information can only be recorded until all such memory units have been excised or flipped, and so the DNA modification rate must be precisely tuned so that the record is not consumed before the lineage of interest is created. It is desireable, then, to create an “unlimited” record so that precise modification rate tuning is not necessary.

The CRISPR system represents a natural chronological record [9] of stimuli where pieces of DNA corresponding to phages are inserted into the genome in the order in which the phages were encountered. Phage genomes are chopped into short oligos, which get inserted into the front of the CRISPR array through the action of the Cas1 and Cas2 proteins. In so doing the CRISPR system can keep inserting more phage sequences and extending the CRISPR array indefinitely, in contrast with more limited integrase-based memory. More recently encountered phages appear closer to the promoter at the front of the CRISPR array, thus those guides are produced in greater abundance than older guides that reside farther down the array. This allows the cell to focus its immune defenses against more pressing threats, while eventually forgetting the faces of long-vanquished foes.

Several groups have endeavored to harness this recording system to create a “DNA tape recorder” circuit. After the function of cas1 cas2 proteins was elucidated [REF], Church et al showed that seeding the cytoplasm with different electroporated oligos[10] Would result in those oligos being preferentially integrated at the beginning of the array. Subsequently, Sheth et al applied the same concept to record the changing copy number of a plasmid, thereby demonstrating the possibility of creating a chronological record of chemical stimuli [11]. In short, overexpression of the spacer acquisition proteins cas1 and cas2 leads to random DNA insertion into the CRISPR array, and by controlling the content of cellular DNA, spacer identity can be influenced. Using this system, the DNA tape recorder made by Sheth et al can identify the presence, absence, and chronological order of three different chemicals over a period of four days. An equivalent “DNA flipping” based event recorder would have to have 81 possible DNA states. Theoretically the system would allow recording to continue indefinitely, but the low rate of spacer acquisition means that eventually the “signal” created by integrating spacers from the plasmid gets diluted out by random DNA.

We sought to design a DNA tape recorder system that was not dependent on the slow and random activity of cas1 cas2 proteins to construct a genetic chronological record of events. Our event logger consists of three components (Fig. 1 A): a set of “data plasmids” that serves as a source of DNA for integration, a synthetic genetic network utilizing phage serine integrases that converts stimulus detection into plasmid integration, and an engineered genomic integration site.

**Figure 1:**
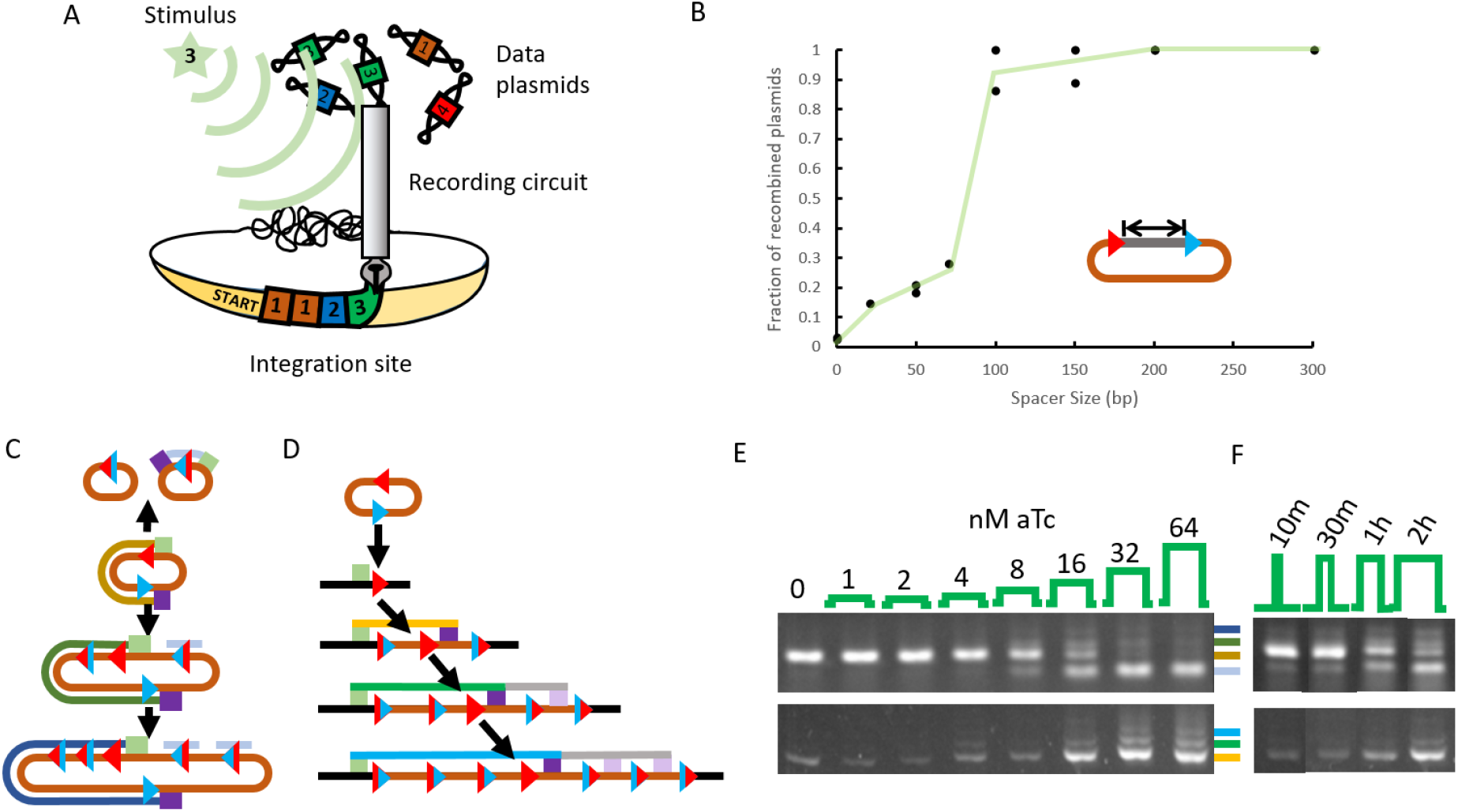
**A**) conceptual representation of the genome event logging circuit. Data plasmids are selected by the recording circuit to be inserted into the integration site, directed by an external stimulus. **B**) Minimal site spacing required for intramolecular integration. Plasmids containing a variable length spacer are incubated with integrase-expressing plasmid in cell-free extract. After incubation for 20 hours, plasmids are purified and transformed into competent cells such that each cell gets one or zero plasmids. Recombined plasmids yield green colonies, while un-recombined plasmids do not express GFP. Colonies were counted and the results are plotted. Green line is purely for visualization. **C**) Possible integrase reactions that affect the data plasmid. Data plasmids can undergo an intramolecular reaction (up arrow) aka deletion, or intermolecular reaction with other data plasmids, aka multimerization. Green and purple squares represent PCR primers used to detect the presence of deleted or multimerized plasmids. Colored lines represent the PCR product that is obtained, with the length increasing from yellow to green to blue. **D**) Data plasmid can integrate into the genome, yielding a repetitive DNA region. Again, PCR primers are designed such that they produce a longer product the more copies of the data plasmid are inserted. Faint violet and grey lines represent copies of the primer sites located further downstream in the genome site. **E**) PCR performed on liquid cultures of bacteria expressing integrase under an inducible promoter. Upon addition of different concentrations of aTc for one hour, data plasmids are recombined to form multimers (top) and the genome site is extended (bottom). Colored lines at the side of the plot indicate which PCR product (depicted in C and D) is seen. **F**) Same as E, but 16 nM of aTc were added for different amounts of time, after which the cells were spun down and resuspended in fresh media without inducer. A similar trend is observed, with more total amount of inducer resulting in more genome integration and more data plasmid multimerization.

We believe that our system offers several advantages over that developed by Sheth et al. First, phage integrases are much more efficient and their recognition site is more well-defined than that of cas1-2, which allows our system to react faster than cas1-2 while being less toxic, since phage integrases will not interact with the *E. coli* genome if their cognate attachment site is not present. Second, our system can allow integration of any size of DNA fragment, which can lead to wider applications such as stimulus-directed pathway assembly or programmed integration of promoters and other active genetic elements. We envision these systems being used to produce “molecular sentinels”—bacteria that can be seeded in a river or a waste treatment plant or a gut microbiome to record chemicals or hormones present over time in a much less obtrusive way than using conventional means.

## Results

Serine integrases will catalyze a recombination reaction between attP and attB sites, converting these into unreactive attL and attR sites [12]. To allow repeated integration into the same site, a data plasmid must contain both attP and attB sites, to replace the attB site which is destroyed by the recombination. This presents a challenge because intramolecular attachment sites may be recombined with much higher efficiency than intermolecular sites, meaning that data plasmids will be consumed in non-constructive reactions faster than they can be integrated into the genome. We determined that placing parallel attachment sites closer than 100 bp, as measured from the edge of the attB and attP sequences, decreases the rate at which intramolecular recombination occurs at a rate inversely proportional to the distance between the sites (Figure 1). The minimum intramolecular integration rate occurs at 0 bp spacing, resulting in less than 5% of the intramolecular integration activity seen with 100bp spacing.

Next, we constructed a proof of concept event logger system consisting of a single data plasmid and tetracycline inducible Bxb1 integrase. Upon induction, integrase was able to catalyze genome integration and data plasmid multimerization in vivo, proportional to the strength of the inducer pulse seen by the cells (Figure 1). Only one hour of induction with aTc concentations ranging from 1-64 nM generated different amounts of integration, with 32 and 64 nM resulting in nearly complete integration (intermolecular or genomic) of all data plasmids. We sought to control the population of data plasmids by arranging the integrase attachment sites in a translational fusion with chloramphenicol resistance. Thus, data plasmids inserted into the genome or multimerized would not be producing a functional antibiotic resistance gene, and the cell would be forced to maintain a population of un-recombined data plasmids to allow for recording future stimuli. However, multimerized data plasmids were not seen to decrease in number even after additional culturing of the cells for 12 hours following the application of the inducer pulse (data not shown). In addition, entire plasmids were integrated into the genome with this system, resulting in multiple functional Cole1 replication origins being present in the genome following integrase induction. We were unable to isolate cells containing different numbers of genome integrated plasmids, possibly because these high copy origins were resulting in polyploidy.

We also wanted to allow recording of the chronological order and duration of multiple events. To this end, the integrase must select between identical attachment sites to integrate different data plasmids into the genome. We have previously shown that catalytically inactive CRISPR-Cas9 could be used to bind and prevent Bxb1 integrase from binding to specific attachment sites in cell-free extract [13]. Now we have shown that this system works in live *E. coli* (Figure 2). A plasmid containing two integrase attachment sites can be made to preferentially integrate one or the other, by co-expression of dCas9 and the appropriate guide RNA. This behavior is dependent on dCas9 expression, but a slight leak of pLac-driven guide RNAs results in colonies nominally expressing only dCas9 to appear similar to dCas9 and second (pLac) guide RNAs in this experiment. As a proof of principle that this system is sensitive to induction time, we also tried activating guide RNA and dCas9 production in advance of integrase production, to see if pre-assembled dcas9-guide RNA complexes were more effective in blocking integrase expression (Figure 2 C). We found that pre-incubation with inducers can increase the magnitude of site selection by about two fold by pre-incubation for 100 min.

**Figure 2:**
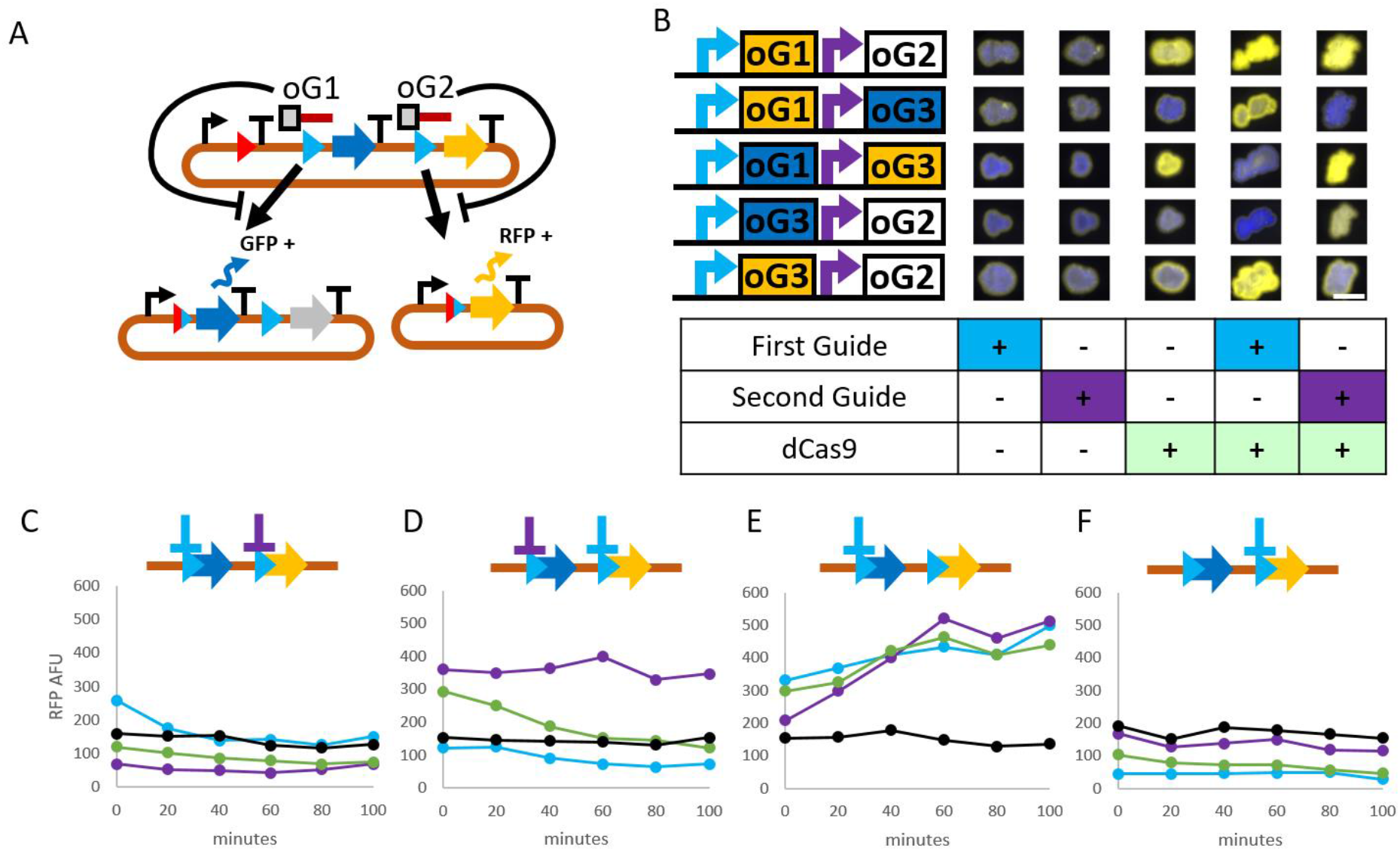
**A**) In vivo integrase attachment site selection reporter construct. Blue triangles are attP sites, the red triangle is an attB site. Bent arrow and T represent promoter and terminator, respectively. In the initial configuration, the promoter is blocked by a terminator and does not transcribe RNA coding for GFP (blue arrow) or RFP (yellow arrow). Upon integrase-mediated recombination with attB and the first attP, the terminator after the promoter is removed and GFP is expressed. Likewise, if the second attP is chosen, the terminator and GFP sequence are removed, allowing RFP to be produced instead. Guide RNAs, represented by grey boxes with orange tails (grey box is at the 3’ end of the gRNA, where the PAM sequence is), can repress integration at either the first or second site. **B**) colonies were produced by spotting cells on agar plates containing different inducers. Constructs with orthogonal guide RNA sequences under control of inducible promoters were induced in the presence of a reporter construct containing guide RNA binding sites. First guide was controlled by an IPTG inducible promoter, and the second guide was controlled by an arabinose inducible promoter. Colored boxes after promoters on the left represent the expected color of the colony if that guide RNA is expressed. For example, the first row has the oG1 binding site in front of GFP, such that if oG1 is expressed, the cells should produce RFP and appear yellow. Consequently, guide binding site for oG3 is present in front of RFP in the first row, so when G2 is expressed, cells should look the same as when no guide is expressed, indicated by a white box. An orange box means that the guide RNA listed inside the box binds to the sequence in front of GFP, such that resulting cells should appear yellow. Scale bar is 5 mm. **C-F**) Guide RNA and dCas9 were induced some amount of time before integrase was induced. Constructs are similar to the ones used in B. Either the first or second guide can be expressed, which are indicated by cyan or purple lines. These in turn lead to repression of the attP site in front of either GFP or RFP on the reporter plasmid. Endpoint RFP fluorescence is plotted. The black line represents cells with no integrase induced, and the green line indicates cells with dcas9 only, which was induced some number of minutes before integrase induction. Cyan and purple lines indicate cells where dcas9 and guide RNA were simultaneously induced before integrase. The first guide RNA was oG1 and the second guide RNA was oG2 **C**) Reporter had the oG1 binding site in front of the GFP attP site, and oG2 binding site in front of the RFP attP site. **D**) Reporter had the oG2 binding site in front of the GFP attP site, and oG1 binding site in front of the RFP attP site. **E**) Reporter had the oG1 binding site in front of the GFP attP site, and oG3 binding site in front of the RFP attP site. **F**) Reporter had the oG3 binding site in front of the GFP attP site, and oG1 binding site in front of the RFP attP site.

We next sought to investigate the feasibility of sequencing as a readout for the memory array content. Since the array will consist of many repeats of known sequences, long error-prone reads should be more than adequate to deduce the identity and order of memory units in the array. We utilized an Oxford nanopore MinION (Oxford, UK) DNA sequencer to sequence PCR products made from genome sites that had data plasmids integrated. For this experiment we utilized a “proof of concept” event logger as described above. In this system, integrase is induced by addition of aTc, and catalyzes data plasmid integration (Fig. 3). Two different data plasmids were tested, but only a single event—that of aTc being present—was recorded.

**Figure 3.**
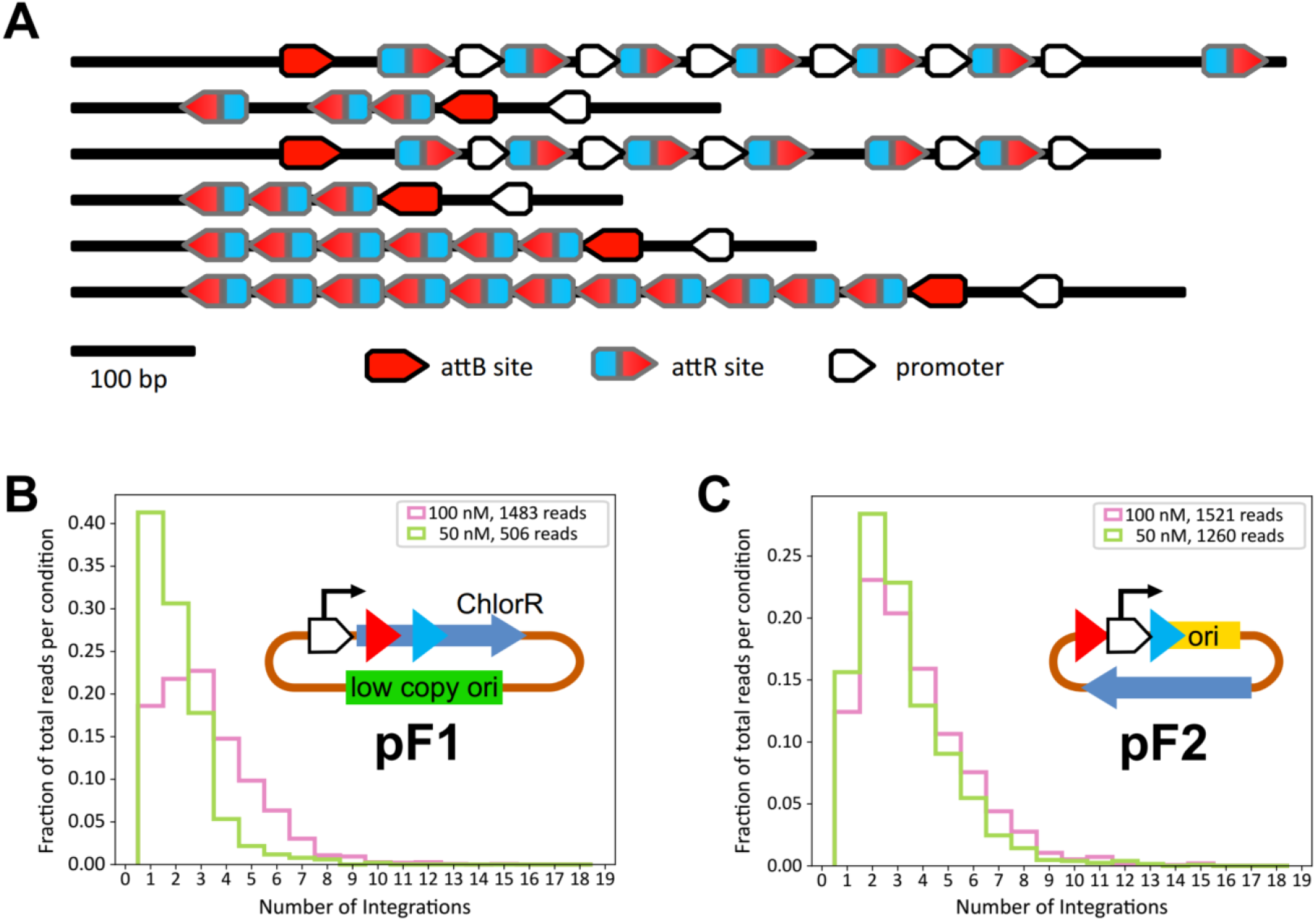
Long read sequencing of multiple integrations. **A)** Representative plot of annotated reads. Each black line represents a single read. Shapes drawn on the line are regions which have > 80% homology to either bxb1 attB, attR, or a constitutive promoter used in the construct. **B)** Cells containing data plasmid pF1 were exposed to 50 or 100 nM aTc, which results in integrase expression from a PTet promoter in the genome. The pF1 plasmid is maintained at low copy number with an SC101 origin of replication, an d has attB and attP sites downstream of a promoter, inside the coding sequence of Chloramphenicol resistance (ChlorR). Average integration site length increased from 2.07 to 3.25 when inducer amount was increased from 50 nM to 100 nM. In this case induction was carried out in minimal M9 media, and inducer concentration was maintained for four hours before cells were lysed and PCRed. **C)** A similar experiment was performed as in B. In this case, pF2 was constructed such that a promoterless Cole1 origin was only capable of plasmid replication before it becomes integrated into the genome. With this plasmid, the distance between attachment sites is greater than in B, yet the average number of integrations observed is approximately similar (3.65 and 3.18 in 100 and 50 nM inducer, respectively).

The first data plasmid (pF1) tested was designed such that chloramphenicol resistance on the data plasmid is abolished once the plasmid is integrated into the genome. This is designed to encourage persistence of the data plasmid after some integration has already occurred, allowing long duration recordings. The pF1 plasmid also contains a low copy SC101 origin, meaning that less data plasmids are present at any given time. When integrase is induced by 50 nM or 100 nM of aTc, on average 2.07 to 3.25 integrations are observed, respectively (Fig 3 B). After integrating, this plasmid no longer produces functional chlor resistance proteins, but it still has a functional origin which probably results in excessive genome replication.

To alleviate the potential genome instability from multiple repeated replication origins present in the same genetic locus, we constructed a second data plasmid (pF2), which utilizes a promoter-less Cole1 origin. Without the native promoter at the front of the Cole1 origin [14], plasmids cannot replicate. An exogenous promoter is provided upstream of the promoter-less origin, flanked by integrase sites. When pF2 integrates into the genome, the exogenous promoter is separated from the promoter-less origin, producing an array of nonfunctional origins in the genomic recording site. This has the advantage of preventing undesired excessive genome replication, but antibiotic resistance is untouched by integration, and after integration has occurred, cells are no longer required to maintain data plasmids. One consequence of the exogenous promoter driving Cole1 origin replication is a greatly increased plasmid copy number[15], since the exogenous promoter is stronger than the native Cole1 promoter. This may be responsible for the increased average integration count at 50 nM aTc of 3.18 when compared to 2.07 with the lower copy pF1 (Fig 3 C).

Each event that this system can record must be represented by a data plasmid containing a unique guide RNA binding site adjacent to and partially overlapping the attP site on that plasmid. Thus we sought to identify the most effective guide RNA location, relative to the bxb1 attP site, that would result in the best blocking while also providing many base pairs of orthogonality. *S. pygoenes* CRISPR-Cas9 binding is sequence specific, relying mostly on the base pairs closest to the Protospacer Adjacent Motif (PAM)[16] thus, it is most critical that the guide RNA sequence closest to the PAM is located outside the integrase attP site, so orthogonal guide sequences are possible. To this end, we designed a series of guide RNAs that overlap the attP site to various degrees (Figure 4 A). In all we tested 8 different guide sequences that bind in the region 5’ of the attP site, 5 that are oriented with the PAM sequence pointing away from the attP site and three with the PAM sequence pointing toward the attP site. We used a very similar construct to the initial integrase site repression construct depicted in figure 2, but this time the guide RNA could only bind to a location adjacent to the attP site upstream of RFP. So if dCas9 can successfully block integrase activity, we should that cultures where guide RNA and dCas9 is induced will produce less RFP and more GFP, corresponding to a bulk shift from choosing the attP site next to RFP to choosing the site next to GFP.

**Figure 4.**
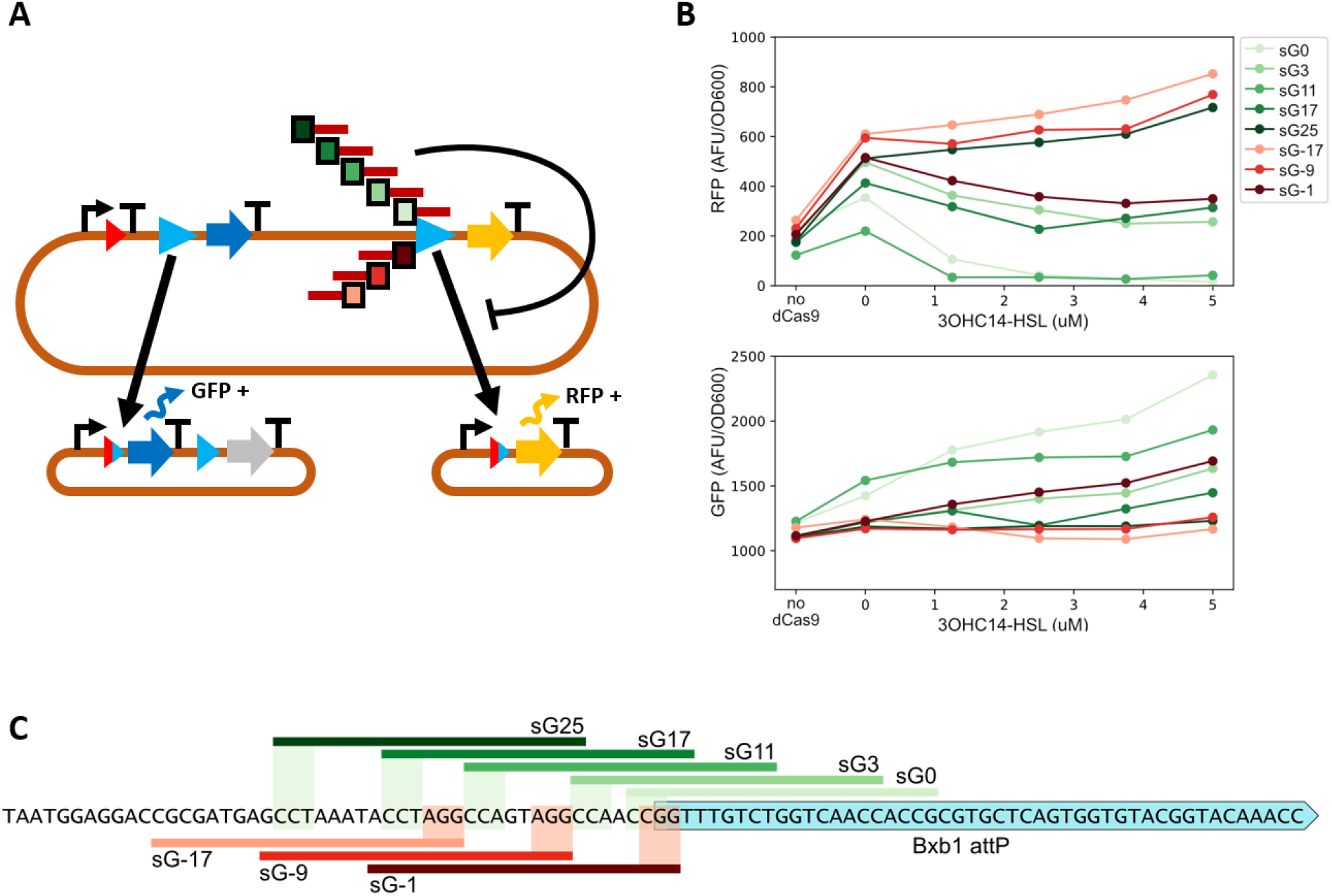
guide RNA spacing optimization. **A)** Construct schematic for spacing optimization experiment. 8 guide RNAs are designed such that they bind at different locations adjacent to the bxb1 attP site (cyan triangle) upstream of RFP (yellow arrow). Guide RNA binding should lead to decreased RFP expression because plasmids that are recombined into the RFP expressing configuration are less numerous if the guide RNA dcas9 complex is in fact interfering with the integrase’s ability to recombine plasmids into the RFP+ state. **B)** endpoint fluorescence values of cells exposed to different amounts of inducer that leads to guide RNA expression. dCas9 is driven by pTet promoter (induced with 300 nM aTc), and integrase by pSal promoter (induced with 100 uM Salicylate). Guide RNA sequences (legend) are depicted in C. **C)** guide RNA sequences used for spacing optimization. Shaded squares outline PAM sequence, colored lines indicate guide sequence. G# denotes the number of nucleotides between the end of the attP site and the 3’-most base pair of the guide RNA. For example, sG3 binds to the PAM sequence CCA (reverse complement is TGG) and so the 3’ most nucleotides of the guide bind to CCA, which is three base pairs outside of the attP site. Negative numbers indicate that the guide binding is on the opposite strand.

Guides that overlap more of the attP site in general have more effect on the integrase activity than guides that are farther away from the attP site. This is demonstrated by the increased GFP and decreased RFP endpoint fluorescence we see when sG0 and sG12 are used, as compared to sG25 and sG-17. Interestingly, sG3 does not follow the pattern that more overlapping guides produce a greater effect, and sG-1, which ostensibly doesn’t overlap at all, produces some effect on integrase activity that seems comparable to sG17. The ability of sG-1 to repress integrase function may be attributable to the steric occlusion of the integrase site by the bulky dcas9 protein, which might take up more space than the guide binding sequence alone. It is unclear why sG3 does not follow the trend that more overlap results in more repression. Perhaps the helical twist of DNA results in the protein-RNA complex being in a conformation that does not impede integrase function. We used different levels of guide RNA induction as a control for leaky expression. Even without adding any inducer, a small difference in GFP and RFP expression is seen when comparing the different guide RNAs tested, which may be a result of leaky guide expression. One would expect that with no dCas9 expression, the integrase should have free reign to choose either attP site, but inexplicably fluorescence of both GFP and RFP are low when dcas9 is not expressed.

We introduced a second data plasmid to the system in order to record the history of two events (Figure 5). Two guide RNAs were used (oG1 and oG2, the same as in Figure 2), such that each guide can repress the integration of one of the two data plasmids. G1 represses the integration of a plasmid containing BC2, so the final record should have a greater number of BC1s when oG1 is expressed, and vice versa for oG2. Cells were induced to express dcas9 and one of the two guide RNAs for one hour, then induced to express integrase. Genome arrays were sequenced and grouped according to how many times BC1 and BC2 were found in each read belonging to a condition. Cells that were made to express guide RNA were compared to cells where no guide RNA inducer was added, to see if there was any change in the composition of the resulting genome arrays.

**Figure 5.**
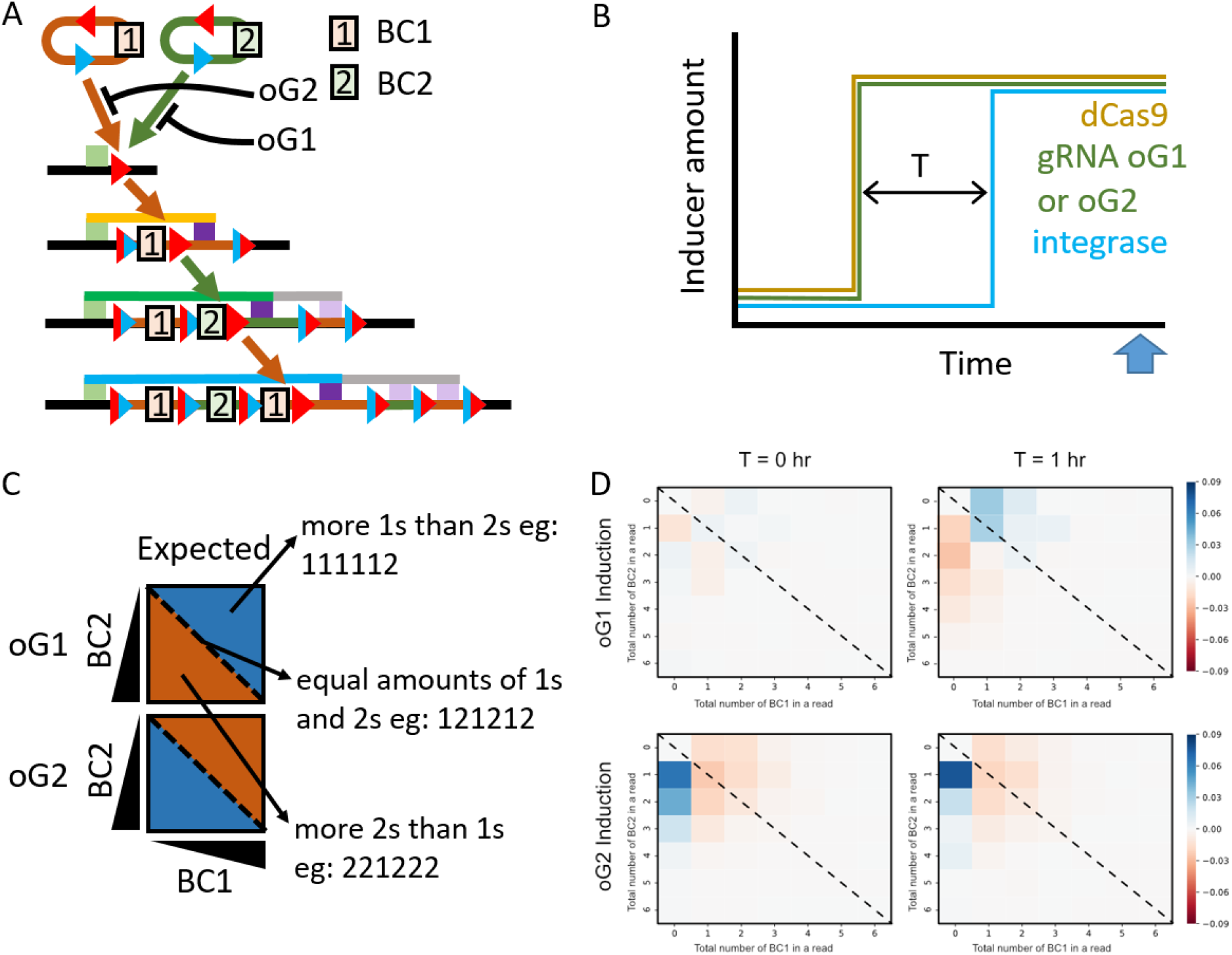
Biased genome array production. **A)** Schematic of the experiment. Two data plasmids are contain barcodes 1 and 2 (square boxes with numbers) and their integration can be blocked by the action of guide RNA 1 and guide RNA 2. Upon integration of either plasmid, the resulting genome array accumulates barcodes in a chronological order, and when proper PCR primers are used (green and purple boxes), a PCR product is made that can be sequenced using Oxford Nanopore MinION technology. **B)** Schematic of time course of inducers added to the recorder cells. First, dcas9 and gRNA are expressed by adding either Salicylate and Arabinose or Salicylate and 3OHC14-HSL. Then, either immediately or after one hour, aTc is added to induce integrase expression. Cells are allowed to grow to saturation and then an endpoint sample is taken and PCRed to prepare for sequencing (blue arrow) **C)** Read content analysis. Reads are analyzed with a 2d histogram where each grid position represents a read composition that contained some number of barcodes 1 or 2. Reads plotted on the diagonal have the same amount of barcodes 1 and 2, whereas reads above the diagonal have more barcode 2s, and reads below the diagonal have more barcode 1s. For example, if G1 has the expected effect of repressing integrase activity, we would see more reads that had more barcodes 1 than 2, resulting in the area below the diagonal to be colored blue in the following heatmap, and vice versa for G2. **D)** Heatmaps of genome array composition. When G1 is induced, we see an increase in reads containing no BC2, but only after one hour of preceding G1 induction. Upon inducing G2, we see an immediate increase in reads containing no BC1. These histograms are generated by subtracting the experimental cells where gRNA was induced from control cells where no gRNA was induced, so the values represent the fraction of total reads that were different from control in each bin. So for example if the value of a bin is 0.03, that means that the experimental culture had 3% more of its reads fall into the bin in question, than the control.

In general the effects on read composition were as expected; induction of oG1 resulted in a decreased number of reads containing BC2 and an increase in the number of reads that contained BC1, and vice versa. Induction of dcas9 and guide RNA before integrase induction improved the magnitude of effects by 3-10 fold, but only in the case of oG1.

## Discussion

We have developed a proof of concept system that allows tape recorder-like sequential recording of stimuli in a bacterium’s DNA. The basic idea is to allow a bacterium to choose between a set of data plasmids to integrate, depending on the stimuli that are perceived. Integrase attachment sites B and P present on the data plasmids allow continued integration of these plasmids into a single B site in the genome, and selective expression of guide RNAs from chemical sensitive promoters will result in binding and repression of attachment site activity, allowing the system to “choose” a data plasmid to insert from a set of possible varieties. In this report we have described a series of steps approaching a complete system capable of genetic recordings, but we are still working to obtain sequences of these repeatedly integrated genome sites. Our goal is to be able to start with a genome site sequence, and then determine the most likely set of stimuli that can create such a sequence. To accomplish this feat, we must first have a system where it is possible to record events, then characterize the system until we understand what a recorded event looks like, and what the limits of event recording are.

In this work we have built a system where event recording is possible, but have not yet determined if it is practical, or what the limits are. To try to understand where we should look for limits or benchmarks on an event recording system it is worthwhile to look at other such systems in the literature. Sheth et al [11] have developed an event recording system that takes advantage of the random spacer integration afforded by cas1-2 protein. By overexpression of this protein, they were able to integrate portions of a “trigger plasmid”, whose copy number is varied by chemical sensitive stimulus. The Sheth et al method shares many similarities to the method proposed here, so it is worthwhile to consider the differences and ponder whether there are tasks more suited to either event recording system.

Both methods involve using a known “data plasmid” that is encoding the event of interest. Sheth et al could easily have multiple such plasmids and control their copy number independently to record events as barcodes just like we propose, and therefore record the same type of events (those that can be translated into transcriptional activation or repression) that we can. The approaches differ in the mechanism of integration and event identification. Using cas1-2 proteins allows the system to record any DNA sequence that is on the data plasmid, so the sequence of the data plasmids is not critical. However, it also results in integration of genomic regions, which could lead to genome damage over time. Our method is much more specific since Bxb1 has no natural binding sites in the e coli genome and this means that we must have very precisely designed data plasmids that contain the sites of interest, at exactly the right spacing. Due to the greater efficiency of Bxb1 we might be able to record events faster, which could lead to greater temporal resolution. Integrases also allow us to append large sequences of DNA, which can be entire genes or promoters. Thus, our system can be used to assemble gene cassettes or pathways in a manner that the cas1-2 integration system cannot.

This brings us to the difference in event detection mechanism. Sheth et al use the copy number of the plasmid to determine which barcode is integrated by the cas1-2 proteins. They have to do this because the cas1-2 proteins are not specific in which piece of DNA they integrate, and by increasing the plasmid copy number the chance that a spacer will come from the plasmid can be controlled. In our system, integrases compete with a dcas9-guide rna complex for binding sites and we can specifically control the frequency with which the integrase binds certain sites. This means our system has a greater potential for orthogonality, since Sheth et al is limited by the number of orthogonal plasmid copy number systems that have been discovered. Because we rely on dcas9-guide RNA binding to detect events, we are subject to the slow assembly and off-rate kinetics of dcas9, which could limit the temporal resolution of events we can record.

At the end of the day the ideal event recorder is different depending on the application. If one desires to record if an event happened without much interest in the specific details of the event, then perhaps a much simpler latching memory system can be used[17]. Recording many different events for long periods of time is at this moment a task best suited to laboratory equipment rather than micro-organisms, so in molecular event recording we must be satisfied with noisy and low time resolution recordings in exchange for dispensing with laboratory equipment.

## Materials and Methods

### Cell strains

Cells used were DH5alpha Z1 from Lutz et al [18]. Genome site constructs were made by Gibson assembly into SpeI-KpnI digested pOSIP KH or pOSIP KO from Pierre et al [19], followed by genome integration and pE-FLP excision protocol as described.

### Constructs

Bxb1 integrase sequence was amplified from the Dual-recombinase-controller vector, which was a gift from Drew Endy (Addgene plasmid # 44456) [6]. dCas9 was amplified from pAN-PTet-dCas9, which was a gift from Christopher Voigt (Addgene plasmid # 62244) [20]. Sequences are listed in table S1.

Recording circuit plasmids were assembled with golden gate followed by Gibson assembly as described [21]. Parts for inducible promoters were modified from plasmids obtained as gifts from Christopher Voigt [22]. pSal, NahR, pTet, pCin, CinR, and pBAD were amplified from source material with primers to add BsaI cut sites using compatible sequences [21] before being used for assembly.

### Experiments

Cells were grown to 0.2 OD in Luria Broth or M9CA minimal media (Teknova) then inducers were added. 1-100 nM Anhydrotetracycline (Sigma), 0.2 uM Sodium Salicylate (Sigma), 0.2% Arabinose (Teknova), 1 mM Isopropyl-beta-D-thiogalactoside (Sigma), or 1-5uM 3OHC14-HSL (Sigma 51481). Cells were subsequently grown for varying amounts of time. Then, 10 uL of cells were transferred into another 1 mL of culture and cells were grown for 12 hours again.

### Sequencing

PCR products for pF1 constructs were obtained using pASS_pF1F: aaaCACCTGCaaaaTTACGGCGTATCACGAGGCAGAAT and pASS_genR: aaaCACCTGCaaaaCCTGGTACAGACAGGAGCTGCGTT. PCR products for pF2 constructs were obtained with pASS_pF2F: aaaCACCTGCaaaaTTACCGGTATCAACAGGGACACC and pASS_genR as above. PCR products were subsequently digested with AarI (NEB), and ligated to barcodes consisting of phosphorylated, dA tailed, annealed oligos having the following sequences:

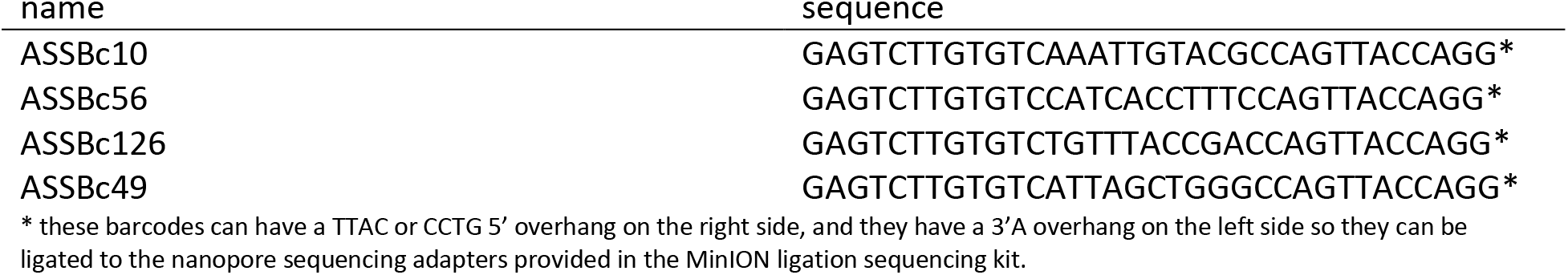

Following barcode ligation, sequencing prep proceeded as described (1D amplicon by ligation (SQK-LSK108) protocol). Loading on MinION R9.4 flow cell proceeded as per manufacturer recommendations.

In other experiments we used primers containing nanopore barcodes such as UintBC1.FOR: AAGAAAGTTGTCGGTGTCTTTGTTGTGGCCTCTGATTGGTGTC and U21BC1.REV: AAGAAAGTTGTCGGTGTCTTTGTTCCGTCTACGAACTCCCAGC. Following PCR with these primers, the DNA was purified and subjected to end repair and ligation as per the SQK-LSK108 protocol.

### Alignment

Sequences were matched to known features using edlib[23] sequencing analysis software. Known feature sequences were as follows:

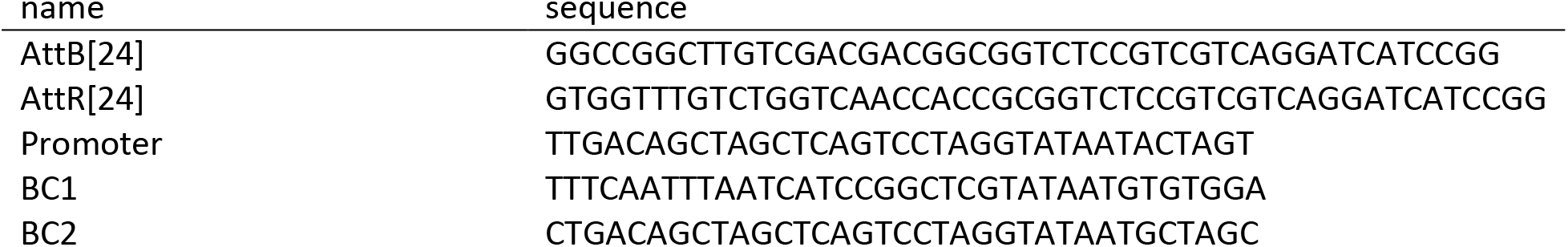

Sequences of constructs used in each figure are reported in supplementary information.

## Supporting information

Constructs

